# Detection of serotonin and serotonin related gene reveals unique roles in human intestinal epithelial development

**DOI:** 10.1101/2025.11.27.690965

**Authors:** Long Phan, Lauren Smith, Kristin M. Milano, Weihong Gu, Madison S. Strine, Harvey J. Kliman, Liza Konnikova

## Abstract

Although 95% of serotonin in the human body is produced in the gastrointestinal tract, most prior research work has focused on the brain. Despite its abundance in the gut, serotonin’s role in early intestinal development remains poorly understood. Here we show that serotonin is detected in both the small intestine and large intestine as early as 14-15 weeks gestational age. Using single-cell transcriptomics with histological validation, we generated a spatiotemporal atlas of serotonin and serotonin-related genes in the human gut from first trimester to late adulthood. The serotonin synthesis gene, *TPH1*, was exclusively expressed in enteroendocrine cells at all time points analyzed, with the highest expression in the second trimester. The serotonin reuptake gene, *SLC6A4*, was largely restricted to mature absorptive enterocytes and increased post-natally. Serotonin receptors also displayed distinct cellular and developmental expression patterns. HTR1E was confined to the first trimester during highly proliferative stage. HTR4 was enriched for in crypt regions, suggesting a role in proliferation for serotonin. Finally, HTR3E was associated with tuft cells throughout development. Together, these findings reveal the dynamic, cell-type-specific regulation of serotonin in the developing human gut, suggesting its importance in early intestinal development, and provide a framework to study its roles in intestinal development and diseases.

## INTRODUCTION

Serotonin (5-hydroxytryptamine or 5-HT) is traditionally regarded as a key neurotransmitter in the central nervous system (CNS) where it regulates mood, cognition and sleep [1]. Interestingly, the majority of serotonin in the human body - around 95% - is produced in the gastrointestinal (GI) tract, predominately by enterochromaffin (EC) cells, a subpopulation of enteroendocrine (EEC) cells [2]. Post-natally, serotonin is also produced by the gut microbiome, which not only synthesizes serotonin directly [3] but also regulates host-derived serotonin production by stimulating EECs [4]. In the gut, serotonin regulates gut motility, modulates immune response, and contributes to gut-brain axis communication [5].

Despite its central role in GI physiology, our understanding of serotonin in the gut remains limited relative to the brain. Most existing works have focused on adult physiology or pathological conditions such as inflammatory bowel syndrome [5], while the role of serotonin during gut development remains underexplored. Increasing evidence suggests that early-life serotonin signaling may influence epithelial maturation or immune interactions [2], which could have long-term effects on GI and systemic health [6].

In the gut, tryptophan hydroxylase 1 (TPH1), the rate limiting enzyme for serotonin synthesis, converts tryptophan to 5-hydroxytryptophan (5-HT), whereas its neuronal counterpart, TPH2, is responsible for serotonin production in the brain [7]. Serotonin can act locallly in the epithelium or be transported to the bloodstream. Serotonin levels are further regulated via reuptake by the serotonin transporter (SERT), encoded by *SLC6A4*. Once transported intracellularly, serotonin is degraded by monomamine oxidase A (MAOA) (**Figure 1A**) [7].

**Figure 1.**
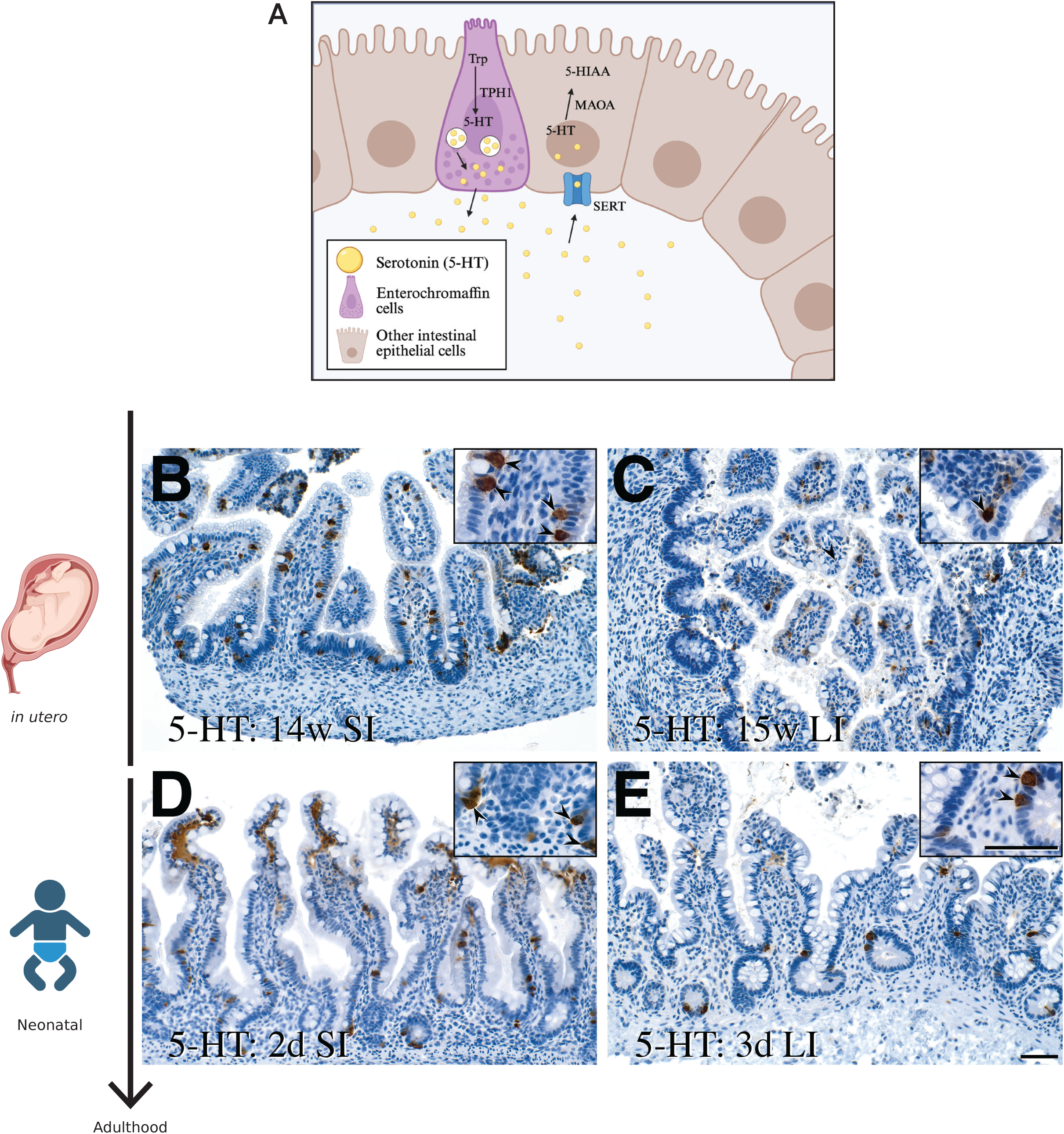
Serotonin expression in early-life intestine. (A) Diagram of serotonin signaling in the gut. (B) Representative immunohistochemistry staining of serotonin (5-HT) in fetal SI tissue, (C) fetal LI tissue, (D) neonatal SI tissue, and (D) neonatal LI tissue. Scale bar: 110 μm.

Serotonin exerts its effects through binding to a diverse family of 5-HT receptors (HTRs), which consists of 15 receptors across seven families (from 5-HT-1 to 5-HT-7) [8]. While most HTRs are G protein-coupled receptors (GPCR) regulating intracellular signaling, the HTR3 family is unique as a ligand-gated ion channel mediating rapid depolarization [9]. Such diversity may allow serotonin to elicit responses that differ in speed, duration and potency. Similar to serotonin itself, the spatiotemporal dynamics of serotonin receptors are better characterized in the CNS. Although several studies have described the expression and functions of HTRs in the gut, emphasis has varied across receptor families, model species, and biological contexts, whether pathological or developmental, resulting in a fragmented view of their roles.

Furthermore, selective serotonin reuptake inhibitors (SSRIs) are widely used during pregnancy. SSRIs can be transferred from the mother to the fetus through placental exchange [10]. Increasing evidence from murine models suggests that exposure to SSRIs can alter fetal brain gene expression [11] and reduce the alpha diversity of the offspring’s gut microbiome [12]. However, the extent to which SSRIs exposure directly affect intestinal development remains poorly understood.

Given the increasing recognition of serotonin’s roles in early-life development [13], addressing these knowledge gaps in the gut’s intrinsic serotonin-producing capacity is particularly important. In this study, we generated a developmental atlas of serotonin and serotonin-related genes across the human lower GI tract, including both small intestine (SI) and large intestine (LI), from as early as 8 weeks estimated gestational age (EGA) to late adulthood. Using single cell transcriptomics (scRNA-seq) and in situ validation with immunohistochemistry (IHC), immunofluorescence (IF) and in situ hybridization (RNAscope), we mapped the presence of serotonin, and expression of serotonin synthesis, transport, degradation, and receptor machinery across cell types and developmental stages.

We found that serotonin can be detected in the small intestine as early as 14 weeks EGA and in the large intestine starting as 15 weeks EGA. Concordant with prior work [14], we observed that serotonin synthesis, reuptake and degradation genes are predominantly enriched in the epithelial compartment of the intestine. The serotonin synthesis gene *TPH1,* although present from 8 weeks EGA, was upregulated in both the LI and SI during the second trimester. Meanwhile, the serotonin transporter *SLC6A4,* although also found prenatally, was enriched primarily in the post-natal SI, highlighting region-specific functions within the developing intestine. Moreover, expression of HTRs also varied across compartments in the gut, including enrichment of *HTR2A* and *2B* in mesenchymal cells, *HTR3A* in B cells, and receptors such as *HTR1E*, *HTR3E* and *HTR4* in the epithelial compartment.

Overall, this atlas provides a framework for understanding serotonergic signaling in intestinal development and may facilitate future studies of how serotonin contributes to gut physiology, the gut-brain axis, and interactions with gut microbiome.

## RESULTS

### Intrinsic serotonin presence in in utero intestinal development

Previous research has demonstrated that maternal serotonin is critical for placenta growth and development [15], highlighting the importance of serotonin in fetal development. However, maternal serotonin represents only one layer of regulation. Using IHC, we identified the presence of serotonin in tissues of the lower GI tract during in utero development (**Figure 1B-E, S1A-I**). IHC staining revealed serotonin expression as early as 14 weeks EGA in the SI (**Figure 1B**) and 15 weeks EGA in the LI (**Figure 1C**). Furthermore, serotonin was consistently detected during gestation through IHC staining in all samples. These data suggested the establishment of intrinsic serotonin production during gestation, where the fetal gut may produce serotonin as early as 14 weeks, or even earlier. As such, we continued to further investigate members of the serotonin production and signaling cascade during human gestation to understand how fetal-derived serotonin production contributes to intestinal development and functions.

### Serotonin synthesis, reuptake, and degradation genes are enriched in the gut epithelium

Since serotonin is a small-molecule metabolite rather than a protein, its presence can not be inferred directly using scRNA-seq. Therefore, we opted to investigate the expression of *TPH1*, *SLC6A4*, and *MAOA* genes to understand the dynamics of serotonin signaling beyond its presence in early life. To do so, we compiled a scRNA-seq atlas of the lower GI tract by combining our previously published SI atlas [16] with additional datasets from the LI [17], appendix [17], rectums [17], and mesenteric lymph nodes [17] (**Supplement Table 1**), yielding approximately 560,000 cells in total (**Table 1**, **Figure 2A, S2A-B**). As predicted, we observed that *TPH1*, *SLC6A4* and *MAOA* were all enriched for in the epithelial compartment of the intestine (**Figure 2B**). Interestingly, *TPH1* expression was restricted to the SI and LI (**Figure S2C**). *SLC6A4* was even more restricted and found mostly in the SI, while *MAOA* was globally expressed in most GI tissues except the intestinal lymph nodes (**Figure S2C**). We then subsetted the UMAP on just the epithelial cells to investigate the expression of these genes within the epithelial compartment (**Figure 2C, S2D**). Upon subsetting, we found that *TPH1* was restricted to a subset of EECs in both the SI and LI (**Figure 2C**). While *SLC6A4* expression was largely restricted to a sub-population of absorptive enterocytes (AE), *MAOA* was expressed across all epithelial sub-types (**Figure 2C**). We further validated that serotonin is produced by a subset of EECs *in utero* using IF, which demonstrated co-localization of serotonin (5-HT) and EEC marker chromogranin A (CHGA). This confirmed that EECs are the primary source of serotonin in the fetal gut, although not all EECs produced serotonin (**Figure 2D**).

**Figure 2.**
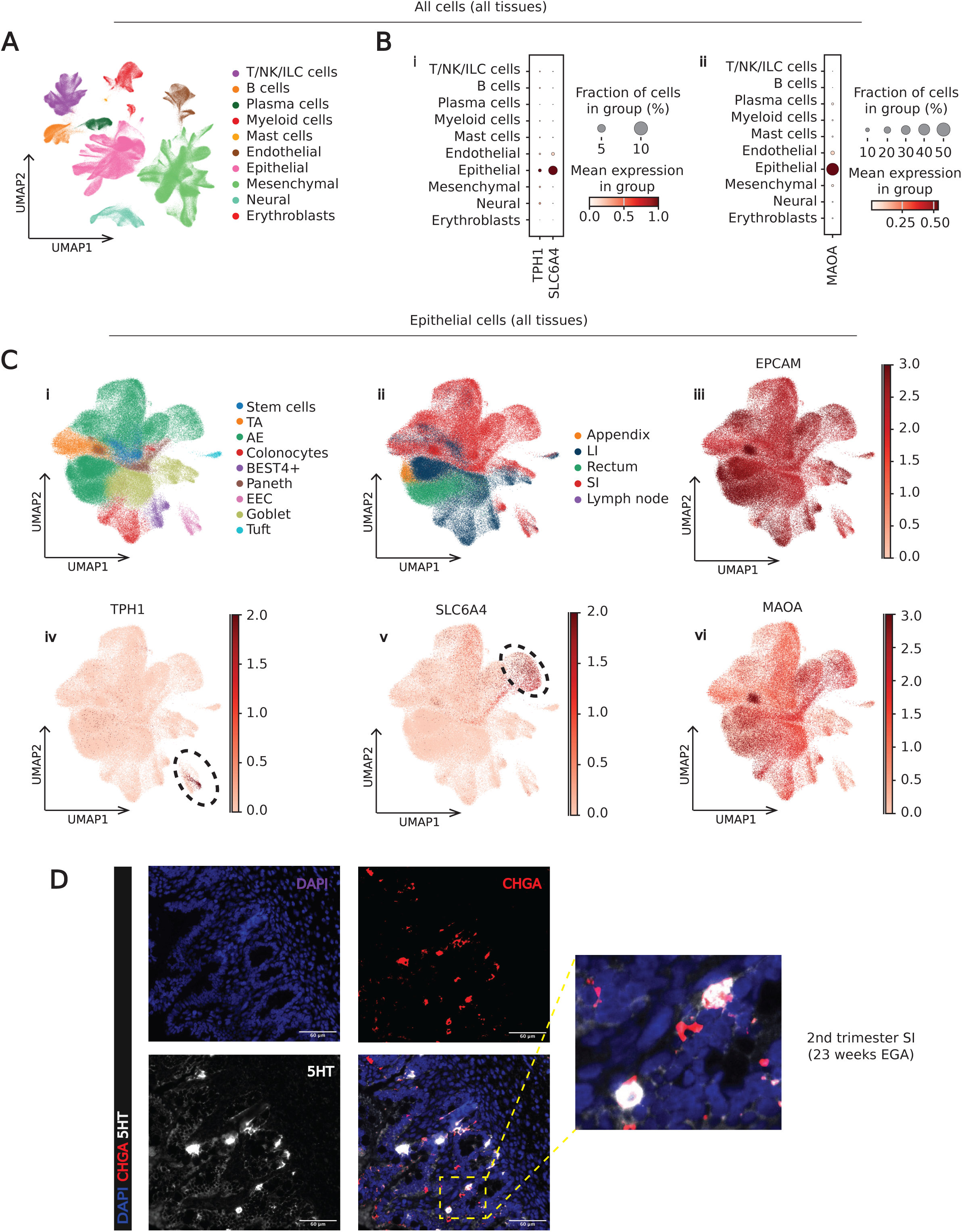
Serotonin synthesis, transportation, degradation genes in the epithelial compartment across tissues and developmental stages. (A) UMAP visualization of the cellular composition of human GI tissues colored by cell types. (B) Dot plot to visualize the expression of (i) TPH1 and SLC6A4 and (ii) MAOA across cell types. (C) UMAP visualization of the composition of the epithelial compartment in all human GI tissues colored by (i) epithelial subtypes, (ii) tissue regions, (iii) EPCAM, (iv) TPH1, (v) SLC6A4, and (vi) MAOA. (D) Representative multiplex immunofluorescence images of enteroendocrine cells (CHGA – red) and serotonin (5HT – white) in fetal SI tissue. Scale bar: 90 μm.

### Cellular restriction of *TPH1* and *SLC6A4* in the small and large intestine

Since not all EEC in the atlas expressed *TPH1*, we then further subsetted the EEC population in the epithelial compartment to identify serotonin-producing subtypes (**Figure 3A-B**) across the SI and LI. Our single-cell data revealed the expression of serotonin-producing EC cells, alongside other EEC subtypes that produce distinct hormones. IHC staining further confirmed the presence of TPH1 in pre- and post-natal GI tissues (**Figure 3C-F, S3A-F**). From our data, we observed that EC cells dominated other EEC subtypes across all developmental ages and tissue regions (**Figure S3G**).

**Figure 3.**
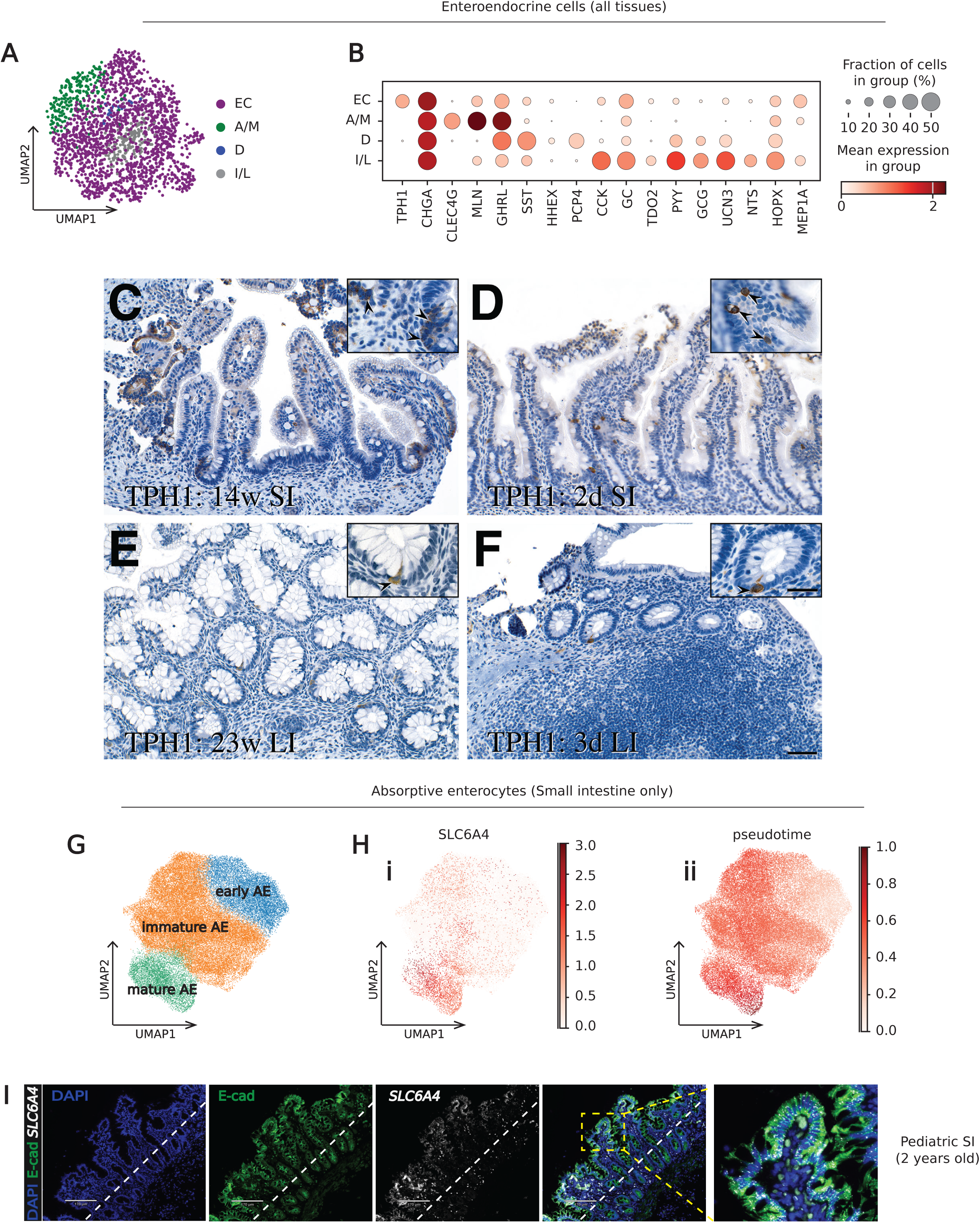
TPH1 and SLC6A4 expression across cell types in the epithelial compartment. (A) UMAP visualization of enteroendocrine subtypes in all GI tissues. (B) Dot plot of marker genes expression in enteroendocrine subtypes across intestines. Color represents normalized mean expression of markers genes in each cell type, and size indicates the proportion of cells expressing marker genes. (C) Representative immunohistochemistry staining of TPH1 in fetal small intestine, (D) neonatal small intestine, (E) fetal large intestine and (F) neonatal large intestine. Scale bar: 50 μm. (G) UMAP visualization of absorptive enterocyte subtypes in the small intestine, colored by subtypes. (H) UMAP visualization of absorptive enterocyte subtypes, colored by (i) SLC6A4 expression, and (ii) diffusion pseudotime score. (I) Representative multiplex in situ hybridization images of SLC6A4 (white) and immunofluorescence for E-cadherin (green) in pediatric SI tissue. Scale bar: 170 μm.

As mentioned earlier, in epithelial datasets spanning all GI tissues, *SLC6A4* was enriched for in a subtype of AEs in the SI. To achieve finer resolution of *SLC6A4* expression, we subsetted the epithelial compartment of the SI (**Figure S3H-I**). This revealed that while *SLC6A4* was detectable at low levels in multiple SI epithelial subtypes, its expression remained consistently enriched for in a subset of AEs, indicating a specialized subpopulation within this compartment (**Figure S3J**). To further characterize this *SLC6A4*-enriched population, we subsetted all SI AEs and grouped them according to their maturation stage along the crypt-villus axis into early, immature, and mature AEs (**Figure 3G, S3K**). Diffusion pseudotime (DPT) [18] analysis demonstrated that AE maturation was associated with increasing *SLC6A4* expression (**Figure 3H, S3L**), with the highest expression in mature AEs. Anatomically, these cells are at the apical end of villi, where AEs are terminally differentiated [19]. This suggests that *SLC6A4* expression is also skewed toward the villus tip of the SI. To validate this, we performed RNAscope for *SLC6A4* combined with IF staining for E-cadherin to visualize *SLC6A4* expression along the crypt-villus axis (**Figure 3I**). Confirming our single-cell transcriptomics findings, *SLC6A4* expression was enriched for towards the villus tip of the SI, suggesting spatial regulation of serotonin in the gut.

### Differential expression of serotonin related genes across development

Importantly, although serotonin-producing EC cells were the major EEC subtypes across tissue regions and developmental ages, this distribution did not necessarily reflect changes in overall *TPH1* expression. The expression levels can vary even within each subset. To capture overall *TPH1* dynamics independent of EC proportions, we examined normalized *TPH1* expression in the epithelial compartment across developmental ages. Using this approach, we investigated whether overall *TPH1* expression changed with age by examining all epithelial cells in the SI (**Figure 4**) and the LI (**Figure S4A-B**). *TPH1* expression was consistently enriched for during the second trimester in both the SI (**Figure 4A**) and LI (**Figure S4C).** We further validated these findings with RNAscope for *TPH1* transcripts combined with IF for E-cadherin (**Figure 4B-C, S4D**). In the SI, we quantified the mean fluorescence intensity (MFI) of *TPH1* expression in all epithelial cells along the crypt-villus axis at three developmental stages: second trimester (23 weeks EGA), pediatric (2 years old), adult (20 years old) (**Figure 4D**), demonstrating that the *TPH1* expression was highest in the second trimester sample, while pediatric and adult samples displayed comparatively much lower but relatively stable expression level.

**Figure 4.**
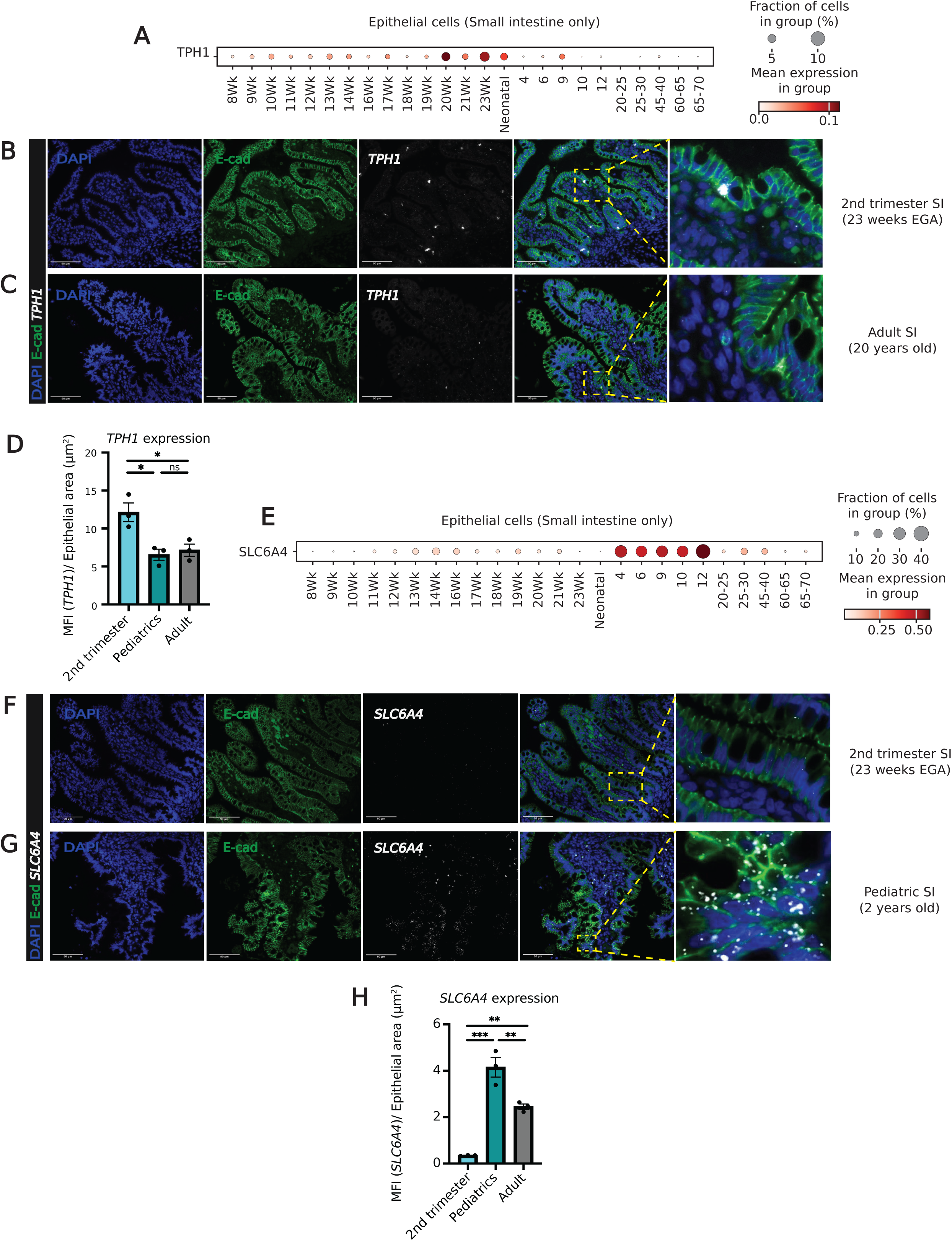
TPH1, SLC6A4, and MAOA expression across developmental ages in the epithelial compartment. (A) Dot plot visualization of TPH1 expression in the epithelial compartment of small intestines across ages. (B) Representative images of fetal second trimester and (C) adult SI tissue. Multiplex in situ hybridization image of TPH1 (white) and immunofluorescence for E-cadherin (green). Scale bar: 90 μm. (D) Mean fluorescence intensity of TPH1 expression in the epithelial compartment along the crypt-villus axis (n = 1 donor per condition, each dot represents a region of interest [ROI], one-way ANOVA, *p<0.05, ns, not significant; data are show as mean ± SEM). (E) Dot plot visualization of SLC6A4 expression in the epithelial compartment of small intestines across ages. (F) Representative images of fetal second trimester and (G) pediatric SI tissue. Multiplex in situ hybridization image of SLC6A4 (white) and immunofluorescence for E-cadherin (green). Scale bar: 90 μm. (H) Mean fluorescence intensity of TPH1 expression in the epithelial compartment along the crypt-villus axis (n = 1 donor per age, each dot represents a ROI, one-way ANOVA, ***p<0.001, **p<0.01; data are show as mean ± SEM)

Next, using the epithelial dataset from the SI, we examined the expression of *SLC6A4* across developmental stages. *SLC6A4* expression was upregulated in pediatric SI samples compared to all age groups (**Figure 4E**), this was further validated with RNAScope/IF (**Figure 4F-G, S4E**). Quantification of *SLC6A4* expression in all epithelial cells along crypt-villus axis at three developmental stages revealed that *SLC6A4* expression was consistently higher in pediatric samples compared to the second trimester and adult samples (**Figure 4H**). Interestingly, the adult sample showed significantly higher *SLC6A4* expression than the second trimester sample, indicating that this expression likely increases post-natally.

When focusing on *MAOA* expression, not only was it ubiquitously expressed in all epithelial subtypes (**Figure S4F**), it was also broadly expressed in the SI (**Figure S4G**) and LI (**Figure S4H**) at all stages assessed, with a slight decrease in late second trimester during the same window as increased *TPH1* expression. However, *MAOA* expression was relatively lower in most secretory lineage epithelial cells, including goblet cells, tuft cells, and EECs, but not in Paneth cells **(Figure S4F)** compared to stem cells, transit amplifying cells and absorptive enterocytes, suggesting some cell type-specific regulation of serotonin degradation within the developing intestinal epithelium.

### Selected serotonin receptors exhibit unique expression in epithelial subtypes throughout the human lifespan

One potential way to restrict the effect of serotonin is through the regulation of its receptors, which we could gain insight into by examining HTRs expression across both spatial and developmental contexts. Our scRNA-seq dataset across all cell types and lower GI tissue demonstrated that serotonin receptor expression was cell type (**Figure S5A**) and developmental stages (**Figure S5B)** specific within the intestines, with overall district patterns. To further investigate these patterns, we examined receptor expression across the intestinal epithelial and mucosal subtypes. The expression of *HTR1* and *HTR6* receptor classes was low in all cell types analyzed, except for *HTR1D*, which was expressed in a small subset of epithelial cells. Notably, both *HTR2A* and *B* were enriched for in mesenchymal cells, while *HTR2B* was also expressed in some endothelial cells. *HTR3A* had the highest expression overall, particularly in neuronal and B cells. *HTR4* was expressed in a subset of epithelial cells and a smaller subset of neuronal cells. Finally, *HTR7* expression was highest in myeloid cells, with notable but weak expression in mesenchymal and neural compartment (**Figure S5A**).

To refine our analysis, we prioritized receptors with clear enrichment in the epithelial compartment - *HTR1D*, *HTR1E*, *HTR3E* and *HTR4* - rather than those predominately expressed elsewhere (**Figure S5C**). Generally, these epithelial-enriched receptors were strongly expressed post-natally and showed higher expression in the SI compared to the LI. Although *HTR1D* was detectable in epithelial cells, its expression pattern resembles that of *HTR4* but with overall lower levels (**Figure S5D-E**). Therefore, we opted to focus subsequent analyses on *HTR1E*, *HTR3E*, and *HTR4*.

While most epithelial-enriched receptors showed postnatal upregulation, *HTR1E* was a notable exception. Within the epithelial compartment, *HTR1E* was primarily expressed in the first trimester in both the SI (**Figure 5A**) and the LI (**Figure S5F**), with expression restricted to approximately 13 weeks EGA or earlier. Within the SI, *HTR1E* was localized specifically to transit-amplifying (TA) cells (**Figure 5B**), whereas in the LI, it was restricted to GPC3+ progenitor cells (**Figure S5G**). Further validation using HTR3E IF staining confirmed HTR1E expression in first trimester GI tissue (**Figure 5C**), while second trimester SI (**Figure 5D**) or adult tissue (**Figure S5H**) showed little to no HTR1E expression. Ki-67 IF staining of the same tissue demonstrated substantial proliferation of intestinal epithelial cells, with some co-localization with HTR1E protein along the forming villous structures in first trimester GI tissue (**Figure 5C-D**). Further MFI quantification confirmed HTR1E expression was restricted to the first trimester (**Figure 5E**).

**Figure 5.**
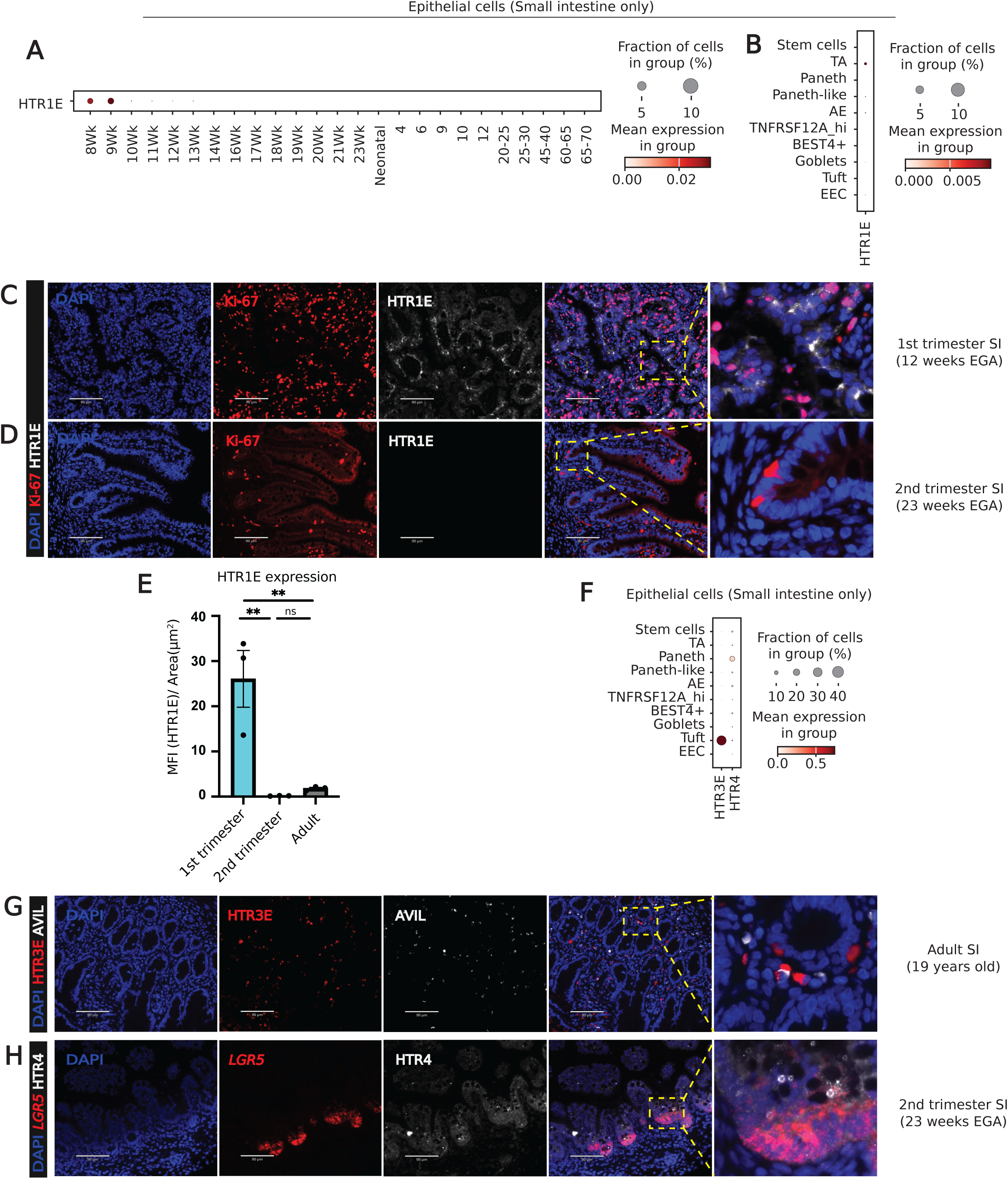
Serotonin receptors HTR1E, HTR3E and HTR4 expression across cell types in the epithelial compartment. (A) Dot plot visualization of HTR1E expression in the epithelial compartment of small intestine across ages. (B) Dot plot visualization of HTR1E expression in the epithelial compartment of small intestine in each cell type. (C) Representative multiplex immunofluorescence images of proliferation marker Ki-67 (red) and HTR1E (white) in first trimester GI tissue and (D) Second trimester SI tissue. Scale bar: 90 μm. (E) Mean fluorescence intensity of HTR1E expression in the intestines (n= 1 donor per age, each dot represents a ROI, one-way ANOVA, **p<0.01, ns, not significant; data are show as mean ± SEM). (F) Dot plot visualization of HTR3E and HTR4 expression in the epithelial compartment of small intestine in each cell type. (G) Representative multiplex immunofluorescence images of HTR3E (red) and tuft cells (AVIL - white) in adult SI tissue. Scale bar: 90 μm. (H) Representative multiplex immunofluorescence images of LGR5+ stem cells (red) and HTR4 (white) in second trimester SI tissue. Scale bar: 90 μm.

In contrast to the temporally restricted pattern of HTR1E, HTR3E exhibited a distinct spatial pattern. We observed that *HTR3E* was uniquely expressed within the tuft cells in both SI (**Figure 5F**) and LI (**Figure S5I**). These patterns suggested an association between serotonin and tuft cell functions, though the directionality of this interaction remains unclear. This was further validated using IF, confirming HTR3E co-localization with tuft cells, marked by Advillin (AVIL), in adult SI tissue (**Figure 5G**). Unlike HTR1E or 3E, *HTR4* expression was much less restricted, and it was present within several epithelial subtypes. Interestingly, *HTR4* was expressed in stem cells and Paneth cells in the SI (**Figure 5F**) and transit-amplifying cells in the LI (**Figure S5J**). Combination of IF with RNAscope staining confirmed the presence of HTR4 within *LGR5+* stem cells (**Figure 5H**).

## DISCUSSION

In this study, we provide a spatiotemporal atlas of serotonin and its associated genes within the human lower gastrointestinal tract from early gestation to late adulthood. Combining single-cell transcriptomics and histological analysis including immunohistochemistry, immunofluorescence and in situ hybridization, we mapped the expression of serotonin, the serotonin synthesis gene *TPH1*, the serotonin re-uptake transporter gene *SLC6A4*, the serotonin degradation gene *MAOA*, as well as selected serotonin receptors - *HTR1E*, *HTR3E* and *HTR4* - across cell types, spatial compartments and developmental stages. Our findings revealed previously understudied cell-type specificity, developmental dynamics, and regional variations of serotonin expression in the human gut.

Our findings extended and confirmed previously established observations that *TPH1* is restricted to EC cells, a subtype of EECs [20]. Consistent with prior reports [21], we identified EC cells as the predominant EECs subtype in both the SI and LI across all developmental ages. Overall, the *TPH1* expression peaked in the second trimester, suggesting heightened serotonin synthesis during mid-gestation. This timing coincides with a critical window of gut development, as the gut undergoes major growth and maturation [22, 23]. Elevated *TPH1* expression likely reflected both an increase in serotonin production during this stage and a potential role for serotonin in epithelial growth and differentiation [7]. This is consistent with previous studies in mouse models [13], which showed that TPH1 expression in the murine gut begins at around embryonic day 15 (E15), roughly equivalent to the human second trimester [24]. The murine work suggested that serotonin present in the gut prior to E15 was unlikely to be produced endogenously, however our study suggests that in humans all the machinery for serotonin production exist early in human gestation. It is likely that in humans there is also a “developmental switch” period in serotonin production where serotonin is initially supplied maternally via the placenta but then produced endogenously by the fetus in fetal gut early in gestation.

Conversely, *SLC6A4* expression, while generally low, was largely postnatal and restricted to absorptive enterocytes in the SI, particularly at the villus tip, highlighting a post-natal shift in serotonin reuptake. Since SERT, the protein encoded by *SLC6A4*, regulates serotonin activities by controlling serotonin availability in the gut [14], its spatial restriction suggests that serotonin’s effects may be more tightly regulated in SI. This pattern is consistent with the classical division of labor in the gut: the SI specializes in nutrient absorption [25], whereas the LI specializes in electrolyte and fluid absorption [26]. Because excessive motility can interfere with nutrient uptake [27], the villus tip localization of *SLC6A4* may represent this fine-tuned mechanism to prevent hyper-serotonergic activity at this absorptive interface. Furthermore, the temporal shift in *SLC6A4* expression may also reflect developmental changes in diet and nutrient needs. The developing fetus acquires nutrients via maternal circulation through the placenta rather than through their GI tract, and neonates primarily consume milk, which is nutrient-dense, pre-digested, and easily absorbed. Meanwhile, children and adults typically consume solid foods, which require direct nutrient extraction and absorption. Accordingly, *SLC6A4* expression peaks in the pediatric phase, reflecting the higher nutrient demands of this stage compared to adults. Simultaneously, these findings imply that the villus-tip enrichment of SERT may serve to clear serotonin quickly and preserve the absorptive niche environment. Beyond their absorptive role, the villus tip is also the most exposed region in the SI, relative to the crypts and lower villus zone. Since the intestinal epithelium forms a critical barrier between the lamia propria and the intestinal lumen, or more broadly, the intestinal immune system and the external environment [28], the villus tip enterocytes constantly undergo shedding [29] and are in constant contact with luminal antigens and microbes [30]. Thus, the villus-tip epithelium also serves as a line of defense, maintaining tight barrier junctions that strictly limit the passage of luminal antigens and macromolecules into the underlying tissue. As serotonin can increase cell permeability and decrease barrier junction integrity [31, 32], excessive serotonin expression in this region may impair the protective role of enterocytes. Given this context, restricting serotonin expression at the villus-tip may help protect the gut from unnecessary downstream immune activation, while supporting efficient nutrient absorption.

Compared to *TPH1* or *SLC6A4*, *MAOA* expression was broadly maintained across epithelial subtypes and developmental stages, reflecting its role as a general mechanism for serotonin degradation [33] and regulation in the gut. Since serotonin, given its small size, can easily diffuse [34] or act non-locally beyond the serotonin-producing cells [35], widespread *MAOA* expression ensures baseline clearance in most epithelial cells. Interestingly, MAOA expression was lower in secretory epithelial cells, including goblet cells, tuft cells, and EECs, which may again reflect functional specializations within the gut. Secretory cells primarily focus on secretion of molecules such as hormones, mucus, or antimicrobial peptides. Meanwhile, absorptive enterocytes are responsible for reuptake, including nutrients and serotonin - consistent with the higher expression of *SLC6A4*, or SERT in these absorptive cells. Serotonin internalized via the serotonin transporter is subsequently degraded by MAOA, which explains the higher expression of *MAOA* in absorptive enterocytes. Moreover, *MAOA* is also highly expressed in crypt regions, including stem cells, TA cells, and Paneth cells. Since MAOA is a mitochondrial protein located on the outer membrane of mitochondria [36], its expression correlates with mitochondrial abundance, which in turn correlates with metabolic activity and the high energy demands within this tissue niche. The overall expression of *MAOA* transiently decreased in late second trimester, coinciding with the rise of *TPH1* expression, suggesting a temporal regulation of serotonin levels. This generally stable expression established a baseline serotonin-clearing capacity, while the transient decrease permitted a critical developmental window of elevated serotonin signaling. Together, these findings highlight dynamic temporal and spatial regulation of serotonin availability in the developing gut epithelium.

Another way to regulate the effects of serotonin beyond degradation or reuptake is through the expression of its receptors. We observed that HTR expression was generally higher in the SI than the LI and tended to increase post-natally. This bias likely reflects SI’s specialization for fine-tuned nutrient absorption. The postnatal increase in receptor expression may reflect developmental transitions in diet and physiology, paralleling with the temporal and spatial shift we observed in *SLC6A4* expression. Thus, such receptor-mediated regulation adds another layer for serotonergic control in the gut, complementing synthesis, reuptake, and degradation.

We found that unlike most serotonin receptors whose expression increased post-natally, *HTR1E* was uniquely restricted to first-trimester epithelial cells, localized to transit-amplifying and progenitor populations in SI and LI respectively. HTR1E is widely regarded as the most uncharacterized serotonin receptor [37], largely because common murine models lack the *Htr1e* gene entirely [38]. Here, we demonstrate that it is uniquely restricted to the first-trimester GI tissues. This developmental window in the GI tract is marked with extensive cellular proliferation, as observed by high Ki67 staining from our data. This observation suggested that HTR1E and therefore serotonin might be acting as a mitogen during this time window [7]. Its restriction to first-trimester tissues also limits accessibility and opportunities for functional study, likely contributing to the relative underrepresentation of this receptor in the literature. Further characterization of the role of HTR1E in early intestinal development would shed light on the potential role of serotonin in early patterning or growth of intestinal tissue.

Furthermore, our data suggest that HTR1E may play a critical role in the formation of stratified epithelial architecture within the intestines. Its restriction to the first trimester may reflect a developmental switch in gut maturation. Since the crypt-villus axis in the SI or colonic crypt in the LI are generally more well-defined after the first trimester [39], the absence of HTR1E expression thereafter is consistent with its transient role. Notably, in mice, crypt-villus zonation emerges post-birth [40], suggesting serotonin signaling through HTR1E may be a human-specific feature of early human gastrointestinal development. HTR1E has also been detected in the brains of other species, such as monkeys and guinea pigs [38]. However, its intestinal expression in non-human primate has not been validated. Further comparative studies will be required to clarify whether this represents a unique human developmental program or a broader mammalian feature.

While there was not a unique temporal pattern to HTR3E expression, it co-localized with tuft cells. Previously, *HTR3E* expression was reported in mouse tuft cells [41], though its function is not well-characterized. Our data demonstrates that *HTR3E* is expressed within tuft cells in both the SI and LI. Its presence in tuft cells across mice and humans suggests that this receptor represents a conserved feature across species. This conservation highlights the potential for studying serotonin-tuft cell interactions in mice and their functional relevance in the human GI tract. In mice, tuft cells are a rare epithelial subtype known for their role in immune sensing and modulation, particularly against parasites and pathogens [42]. In humans, however, their intestinal functions remain more enigmatic [17, 43]. Notably, previous work has shown that mouse tuft cells can take up or respond to serotonin from neighboring enterochromaffin cells, suggesting a serotonin-tuft cell axis of communication [44]. This co-localization may suggest a potential role for serotonin in immune modulation and indicates a conserved interaction in serotonin-tuft cell axis between mouse and human, although tuft cells may perform distinct roles across tissues and species.

In addition, we showed HTR4 co-localized with stem cells and expressed in Paneth cells in SI. Stem cells are essential for epithelial proliferation [45]. Meanwhile, Paneth cells secrete antimicrobial peptides [46], and structurally and metabolically support optimal stem cell function [47]. This co-localization suggests a potential role in serotonin signaling in the crypt area of the intestine, particularly in epithelial development and antimicrobial immunity, which remains poorly characterized. This expression further reinforces the role of serotonin in the crypt rather than villous region, aligning with our data on the expression pattern of SLC6A4, which limits serotonin signaling in the villus tip.

Our understanding of serotonin functions has increased in recent years. As a key modulator in both the gut and the brain, serotonin plays a central role in gut-brain communication. Together, this atlas serves as a valuable resource for understanding serotonin spatiotemporal expression, providing a robust framework to understand early gut development, interactions with gut microbiota, and impacts on immune compartment. The spatiotemporal maps we present could also facilitate studies of drug exposure, such as SSRI use during pregnancy, or investigations of early life disruptions that contribute to gastrointestinal or metabolic disorders. By defining the spatiotemporal pattern of serotonin, serotonin synthesis, transportation and degradation genes, along with the serotonin receptors in the gut epithelium, this atlas further provides critical baseline to interpret how perturbations - genetics, microbes, pharmacologics - may contribute to disorders such as intestinal bowel syndromes, colorectal cancers or metabolic diseases. While our study offers a robust descriptive atlas, our data interpretation and validation are generally restricted to the epithelial compartment, while serotonin receptor expression, especially in other compartments such as immune, remains largely unexplored. This study, while comprehensive in its coverage of large and small intestinal data across developmental stages, is more limited in other tissue regions. Functional validation of serotonin signaling in specific epithelial populations will also be informative and represents an important next step. Moving forward, exploring receptor-mediated signaling pathways in the context of gut development or disease will further enhance the utility of this atlas. Furthermore, we also noted some caveats in our HTR1E data. Our atlas was developed by repurposing previously published single-cell datasets from multiple studies, which may introduce potential uncertainties in regional annotation during the first semester. At this stage, the differences in SI and LI are not as well defined. While we adhered to the original authors’ tissue region designation, we remain cautious about their accuracy at this developmental stage. For our in-house validation staining using first-trimester tissue, we therefore labeled samples broadly as GI rather than LI or SI. Consequently, while we are confident in the temporal-specificity of the first-trimester data, cell-type specificity is less certain and may be influenced by post-first-trimester datasets, where intestinal regions are more clearly defined.

In conclusion, this work provides a spatiotemporally resolved, cell-type-specific map of serotonin signaling in the human lower gut, spanning from early gestation to late adulthood, accompanied by orthogonal validation using IHC, IF, and RNAscope. This emphasis of this atlas in early-life gut development also defines several poorly understood trends in serotonin signaling along the epithelial development trajectory. Ultimately, this atlas lays the foundation for future mechanistic studies of serotonergic regulation in intestinal development and provides a valuable reference for the broader scientific community.

## EXPERIMENTAL MODEL AND STUDY PARTICIPANT DETAILS

### METHODS

#### Sex as a biological variable

Each experiment was conducted independent of sex. Therefore, the data represented in this study is reported as combined male and female.

#### Single-cell dissociation and sequencing

The single cells were isolated from fresh tissues or cryopreserved Biopsies as previously reported. Briefly, tissues were digested using Liberase [17, 48], Neural Tissue Dissociation Kit [49, 50], dispase [51], or collagenase [52, 53, 54]. Some dissociated cell suspensions were further enriched by magnetic cell separation (Miltenyi MACS) with EPCAM positive selection [48] or CD45 positive and negative selection [17, 53, 54]. Sample libraries were prepared using the Chromium 10x Genomics single-cell library kit with 3’ version [48, 49, 50, 51, 52, 53, 54] or 5’ version [17]. Libraries were then sequenced on an Illumina Hi-seq 4000 [17, 48, 52, 53] or Illumina Hi-seq 2500 [49, 50].

#### Analysis of single-cell data

Single-cell transcript counts output from Cell Ranger (10x Genomics) for fetal and postnatal samples were obtained from previously published datasets (E-MTAB-8901, E-MTAB-8901 [48], E-MTAB-9543, E-MTAB-9533, E-MTAB-9536[17], E-MTAB-10187[49], E-MTAB-9363 [50], E-MTAB-9489 [51], GSE178088 [52], https://doi.org/10.5281/zenodo.5457926 [53], https://doi.org/10.5281/zenodo.7780794 [54]) (Supplement Table 1). Clustering analysis was performed as previously described [16]. Briefly, Scanpy (v1.11.2) [55] package was applied to the scRNA-seq data for preprocessing and clustering. Cells that have less than 200 genes and high mitochondrial reads were filtered, and genes that express in less than 3 cells were filtered. Scrublet (v0.2.3) [56] was used to detect and remove doublets, with doublet score produced. Gene expression for each cell was normalized and log-transformed. Following preprocessing, highly variable genes were identified using the scanpy.pp.highly_variable_genes function with default parameters. In addition, the effects of mitochondrial gene percentage, ribosomal gene percentage, and unique molecular identifier (UMI) counts were regressed out using scanpy.pp.regress_out function before scaling the data.

Batch correction of samples was performed using bbknn (v1.5.1) [57] on 50 principal components (PCA). Dimensionality reduction and Leiden clustering were carried out on the remaining highly variable genes, and the cells were visualized using Uniform Manifold Approximation and Projection (UMAP) plots. The clusters with the highest predicted doublet scores were excluded from the analysis. Cell lineages were manually annotated based on known markers genes found in the literature, in combination with algorithmically defined DEGs in each cluster, using scanpy.tl.rank_genes_groups function with the Wilcoxon test.

#### Diffusion pseudotime analysis of absorptive enterocytes

Pseudotime trajectories were inferred using DPT [18] on Scanpy in Python. Briefly, a neighborhood graph was first computed using the top 50 PCA and 15 nearest neighbors. Diffusion maps were then calculated to capture the major axes of variation in the dataset. To root the trajectory, we selected the representative cells from the early absorptive enterocyte cell population, which we defined based on marker gene expression and cluster annotation. DPT was then computed using the rooted diffusion map, yielding a continuous trajectory score for each cell. Pseudotime values were visualized on UMAP and boxplot to examine developmental trajectories across absorptive enterocyte population in small intestines.

#### IF staining

Formalin-fixed, paraffin-embedded (FFPE) intestinal tissues were sectioned at 4-5 μm thickness sections for staining. FFPE slides were embedded and sectioned at Yale Histology Core. Slides were deparaffinized using xylene and alcohol and placed in 1X citrate buffer at 95% for 45 minutes. Slides were then washed in distilled H2O (ddH_2_O) and 1X Dulbecco’s phosphate buffered solution (DPBS). Slides were blocked with 10% horse serum (HS) in DPBS and incubated with primary antibodies diluted in 2% HS in DPBS. After washing, slides were then incubated with secondary antibody in 2% HS in DPBS.

#### Multiplex IF with Tyramine signal amplification

After antigen retrieval, slides were treated with 3% H_2_O_2_ in ddH_2_O. Slides were then blocked in 10% HS as described above and incubated overnight with primary antibody at 4°C. After washing, slides were incubated with HRP-conjugated anti-rabbit secondary antibody for one hour. Tyramide reagent was diluted with 0.15% H_2_O_2_ and Tris-HCl (pH 7.4) buffer in 1:1:100 ratio. Slides were then incubated with the tyramide reagent dilution for 10 minutes prior to development. While submerging slides in 1X citrate buffer solution, slides were microwaved at 100% power for 1 minute and 20% power for 15 minutes to quench the tyramide reaction. Slides were then re-blocked with 10% HS solution before undergoing another round of normal IF incubation.

#### IHC staining

IHC staining was performed on five-micron serial sections using EnVision+ HRP Rabbit DAB+ as previously described [58, 59].

#### In situ hybridization (RNAscope) combined with IF

RNAscope was performed on FFPE slides using the RNAscope Multiplex Fluorescent Reagent Kit v2 with TSA Vivid Dyes assays (Advanced Cell Diagnostics) according to the manufacturer’s instructions. In brief, paraffin sections were deparaffinized and treated with citrate buffer. Slides were then hybridized sequentially with target probes. After RNA signal development, a blocking step was performed with 10% HS and 1% bovine serum albumin in Tris-Buffered Saline, followed by overnight incubation at 4°C with desired primary antibody. Slides were then further incubated in secondary antibody as previously described.

#### Imaging and quantification

Images were taken at 10x and 20x magnification using the Echo Revolve microscope. RNA signals appear as dots. Mean fluorescence intensity was quantified using Fiji [60].

#### Statistical analysis

Statistical significance was analyzed using ANOVA in Prism (v10.0.0). For all comparisons, one-way ANOVA was applied, followed by Tukey’s post-hoc test. Details of the statistical tests used, along with corresponding p values, means, medians, and SEM, are provided in the figures and figure legends. Data are shown as mean ± SEM. p < 0.05 was considered statistically significant.

#### Study Approval

We combined data collected from human intestinal tissues obtained from patients undergoing elective termination of pregnancy or surgeries with IRB approval and informed consent at University of Pittsburgh or Yale School of Medicine (IRB #2000028360). First trimester tissues were obtained from the Research Centre for Women’s and Infant’s Health BioBank in Toronto, Canada under Material Transfer Agreement.

#### Data availability

All data associated with this study are included in the paper and supplementary data. Major code used in this study is available on GitHub (https://github.com/LDPhan/serotonin_atlas)

## Author Contributions

Conceptualization: LK, HK

Methodology: LP, LS, KM, WG, MS, LK, HK

Investigation: LP, LS, WG

Visualization: LP, LS, KM, HK

Funding acquisition: LK, HK

Supervision: LK

Writing – original draft: LP

Writing – review & editing: LP, LS, KM, WG, MS, HK, LK

## Acknowledgements

We thank Serrena Singh from Yimlamai Laboratory, Kerri St Denis, Oluwabunmi Olaloye from Konnikova Laboratory, and Robert Coweles for their constructive feedback and helpful discussions.

---

**Figure S1. Serotonin expression in early-life intestine (cont.)** (A-E) Representative immunohistochemistry staining of serotonin (5HT) in fetal SI tissues, (F) neonatal SI tissue, (G-H) fetal LI tissues, (I) neonatal LI tissue, and (J) pediatrics LI tissue. Scale bar: 110 μm.

**Figure S2. Serotonin synthesis, transportation, degradation genes in the epithelial compartment across tissues and developmental stages (cont.)** (A) Dot plot of marker genes expression in cell types across intestines. Color represents normalized mean expression of markers genes in each cell type, and size indicates the proportion of cells expressing marker genes. (B) Proportion bar plot of cell types in each region. (C) Dot plot to visualize the expression of (i) TPH1 and SLC6A4 and (ii) MAOA across tissue regions. (D) Dot plot of marker genes expression in epithelial subtype types across intestines. Color represents normalized mean expression of markers genes in each cell type, and size indicates the proportion of cells expressing marker genes.

**Figure S3. TPH1 and SLC6A4 expression across cell types in the epithelial compartment (cont.)** (A-C) Representative immunohistochemistry staining of TPH1 in fetal SI tissues, (D) neonatal SI tissue, (E) neonatal LI tissue, and (F) pediatrics LI tissue. Scale bar: 110 μm. (G) Proportion bar plot of enteroendocrine subtypes in (i) each region and (ii) age group. (H) UMAP visualization of epithelial compartment of the small intestine only, colored by subtypes. (I) Dot plot of marker genes expression in epithelial subtype types across small intestine only. Color represents normalized mean expression of markers genes in each cell type, and size indicates the proportion of cells expressing marker genes. (J) UMAP visualization of epithelial compartment of the small intestine only, colored by SLC6A4 expression. (K) Dot plot of marker genes expression in absorptive enterocytes subtypes across small intestine only. Color represents normalized mean expression of markers genes in each cell type, and size indicates the proportion of cells expressing marker genes. (L) Box plot distribution of diffusion pseudotime values for each absorptive enterocyte subtype. The central line indicates the median pseudotime, the box spans the interquartile range (IQR), and whisker extends to 1.5xIQR. Outliers are shown as individual points.

**Figure S4. TPH1, SLC6A4, and MAOA expression across developmental ages in the epithelial compartment (cont.)** (A) UMAP visualization of epithelial compartment of the large intestine only, colored by subtypes. (B) Dot plot of marker genes expression in epithelial subtype types across large intestine only. Color represents normalized mean expression of markers genes in each cell type, and size indicates the proportion of cells expressing marker genes. (C) Dot plot visualization of TPH1 expression in the epithelial compartment of large intestines across ages. (D) Representative images of pediatric SI tissue. Multiplex in situ hybridization image of TPH1 (white) and immunofluorescence for E-cadherin (green). Scale bar: 90 μm. (E) Representative images of adult SI tissue. Multiplex in situ hybridization image of SLC6A4 (white) and immunofluorescence for E-cadherin (green). Scale bar: 90 μm. (F) Dot plot visualization of MAOA expression in the epithelial compartment across epithelial subtypes of (i) small intestine and (ii) large intestine. (H) Dot plot visualization of MAOA expression in the epithelial compartment of small intestines across ages. (H) Dot plot visualization of MAOA expression in the epithelial compartment of large intestines across ages.

**Figure S5. Serotonin receptors HTR1E, HTR3E and HTR4 expression across cell types in the epithelial compartment (cont.)** (A) Dot plot visualization of serotonin receptor gene expression across cell types. (B) Dot plot visualization of serotonin receptor gene expression across all cell types by tissue regions and developmental stages. (C) Dot plot visualization of serotonin receptor gene expression across all epithelial cells by tissue regions and developmental stages. (D) Dot plot visualization of HTR1D expression in the epithelial compartment of small intestines across epithelial subtypes. (E) Dot plot visualization of HTR1D expression in the epithelial compartment of large intestines across epithelial subtypes. (F) Dot plot visualization of HTR1E expression in the epithelial compartment of large intestines across ages. (G) Dot plot visualization of HTR1E expression in the epithelial compartment of large intestines across epithelial subtypes. (H) Representative multiplex immunofluorescence images of proliferation marker Ki-67 (red) and HTR1E (white) in adult SI tissue. Scale bar: 90 μm. (I) Dot plot visualization of HTR3E expression in the epithelial compartment of large intestines across epithelial subtypes. (J) Dot plot visualization of HTR4 expression in the epithelial compartment of large intestines across epithelial subtypes.

## Supplements

**Supplement Table 1. Tissues metadata**

**Supplement Table 2. Antibodies and reagent**

