## Supplementary figures and images for "Detection of serotonin and serotonin related gene reveals unique roles in human intestinal epithelial development"

### Figure S1-3.pdf

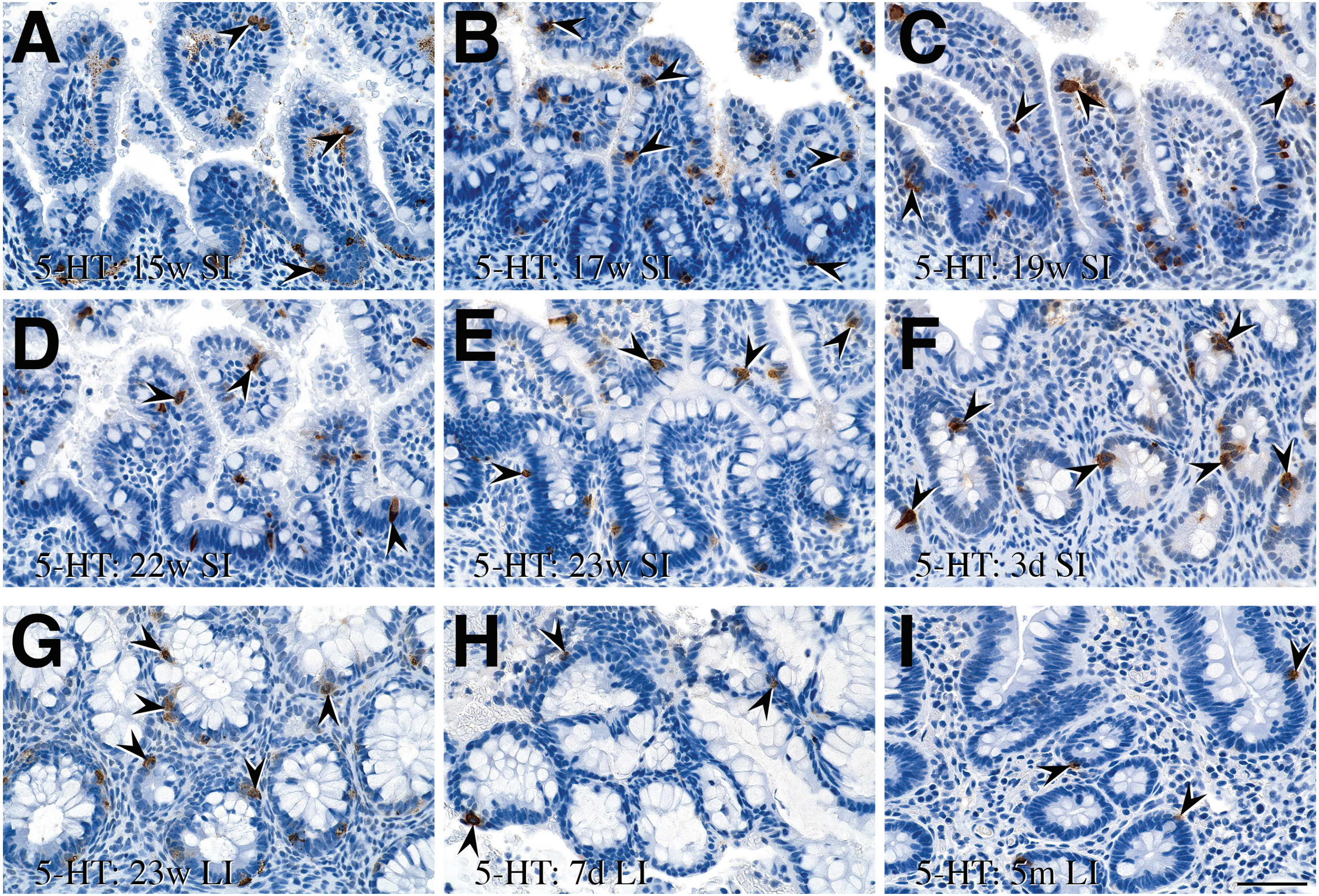

### Figure S2.pdf

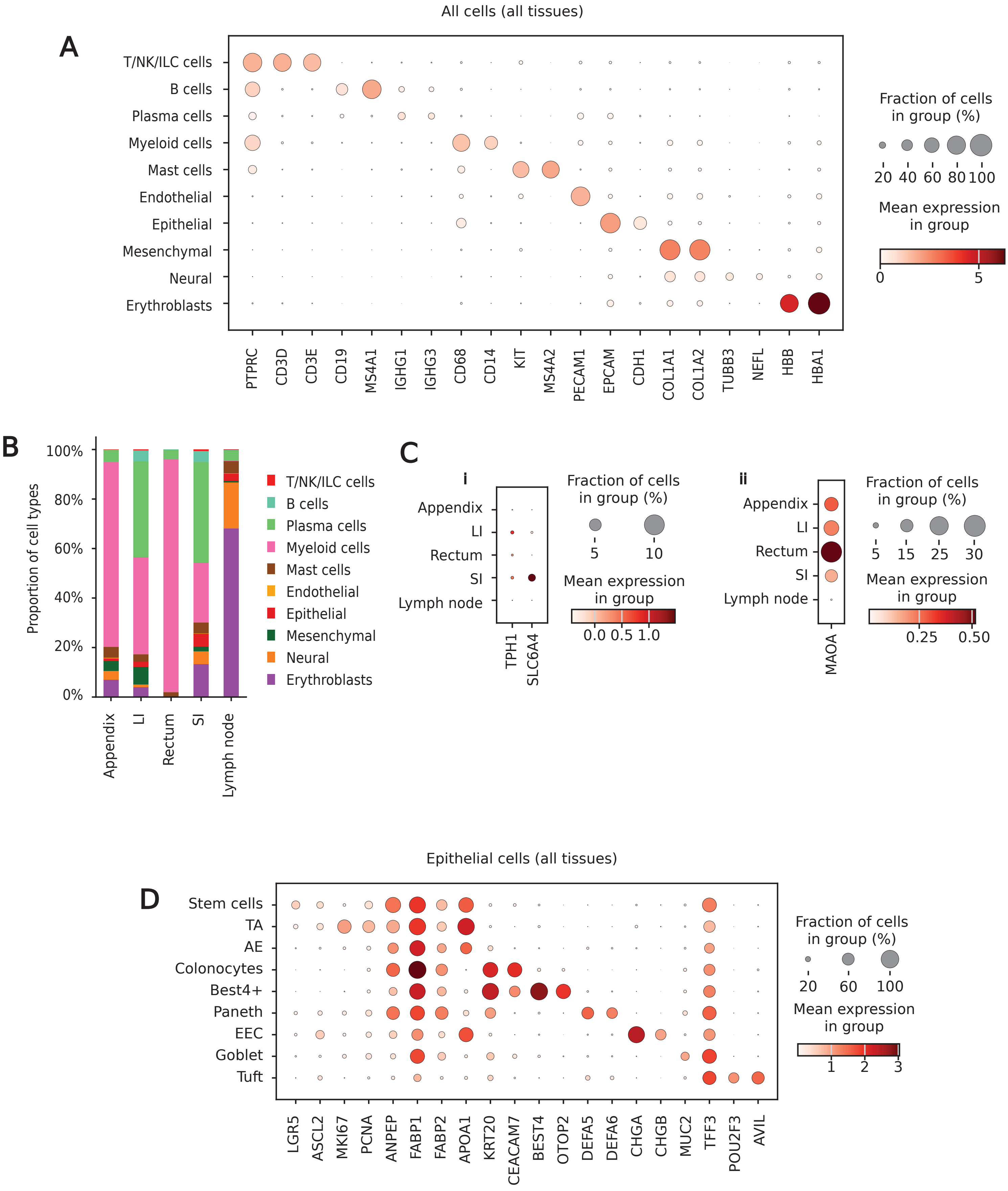

### Figure S3.pdf

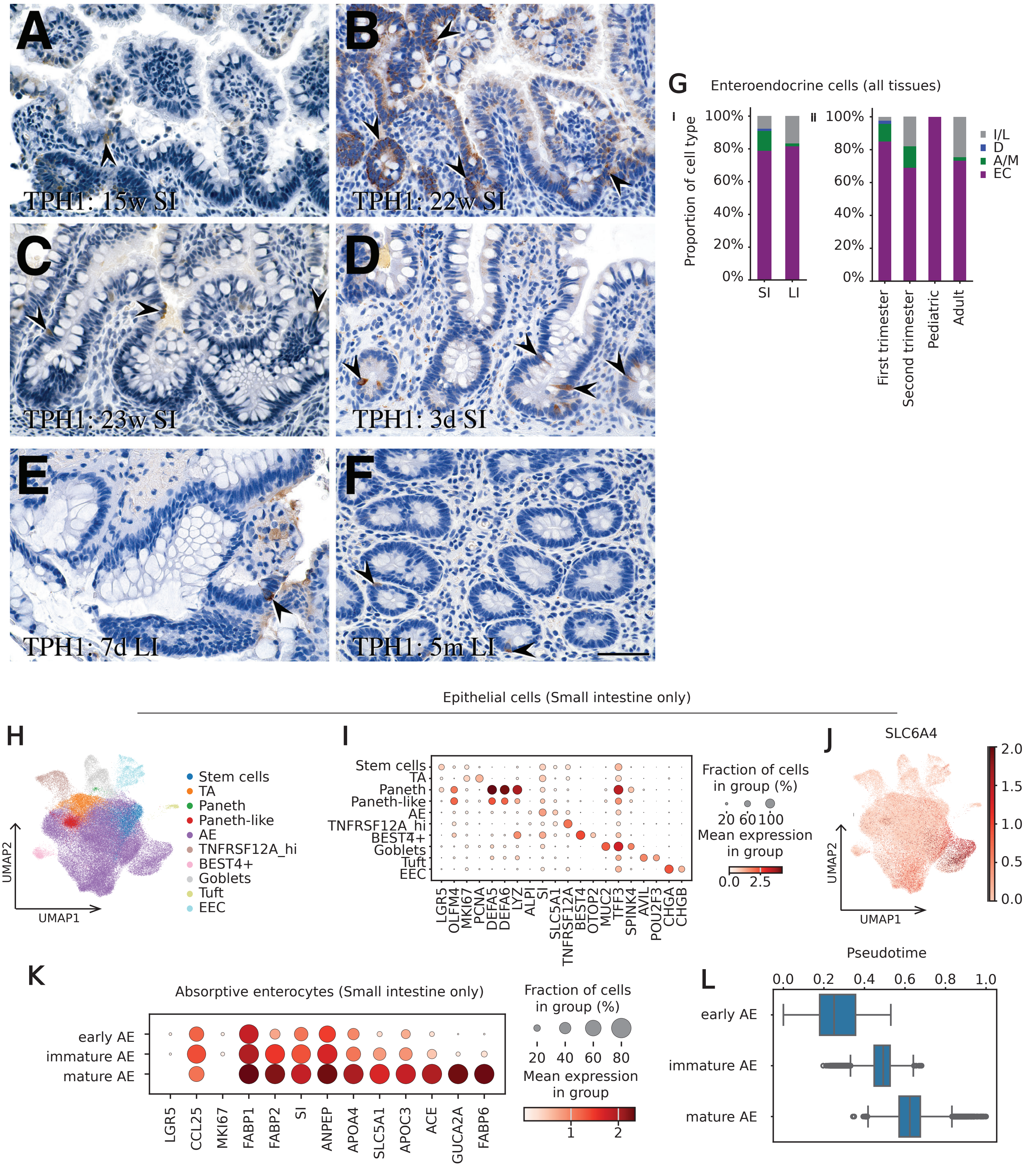

### Figure S4.pdf

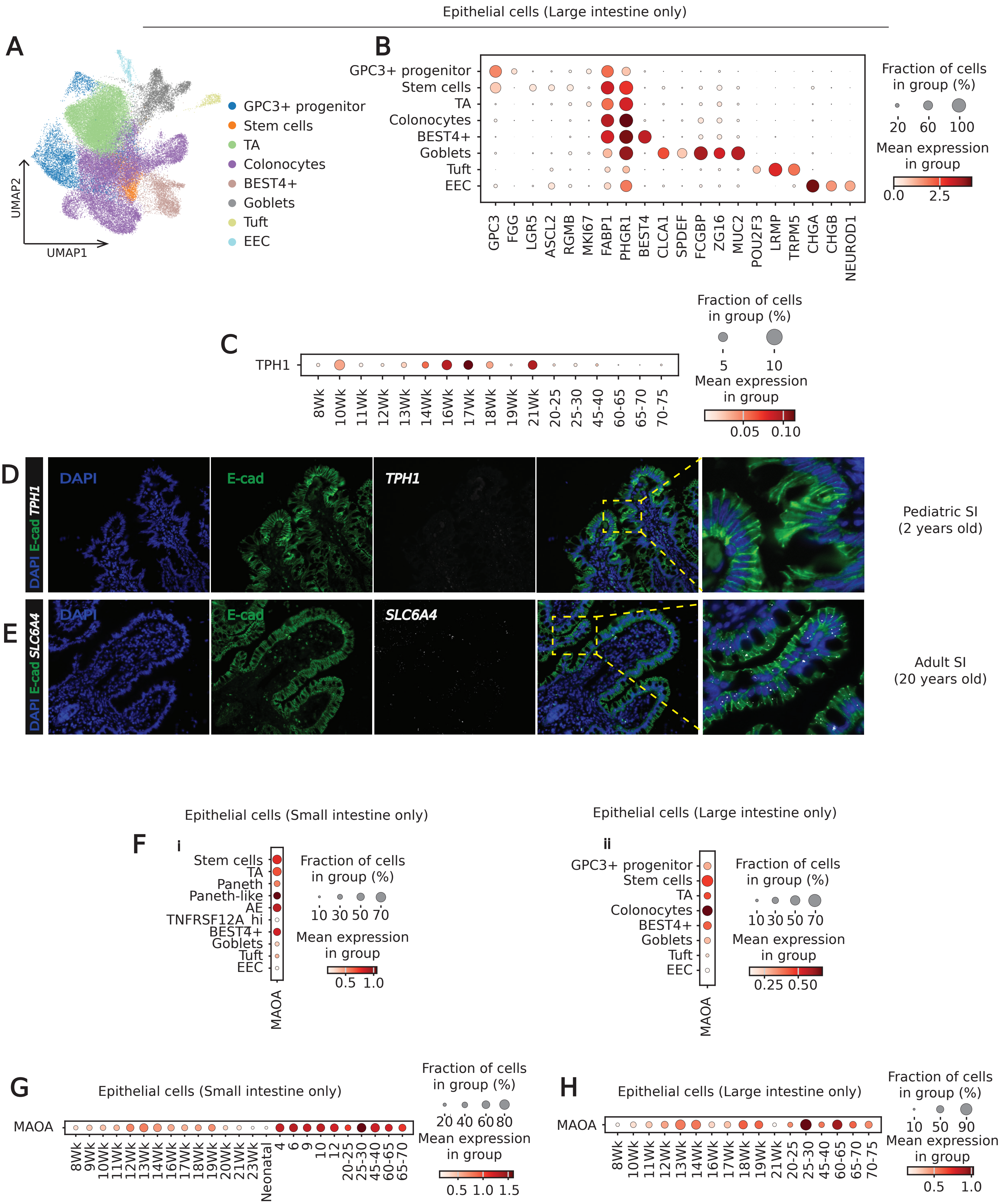

### Figure S5.pdf

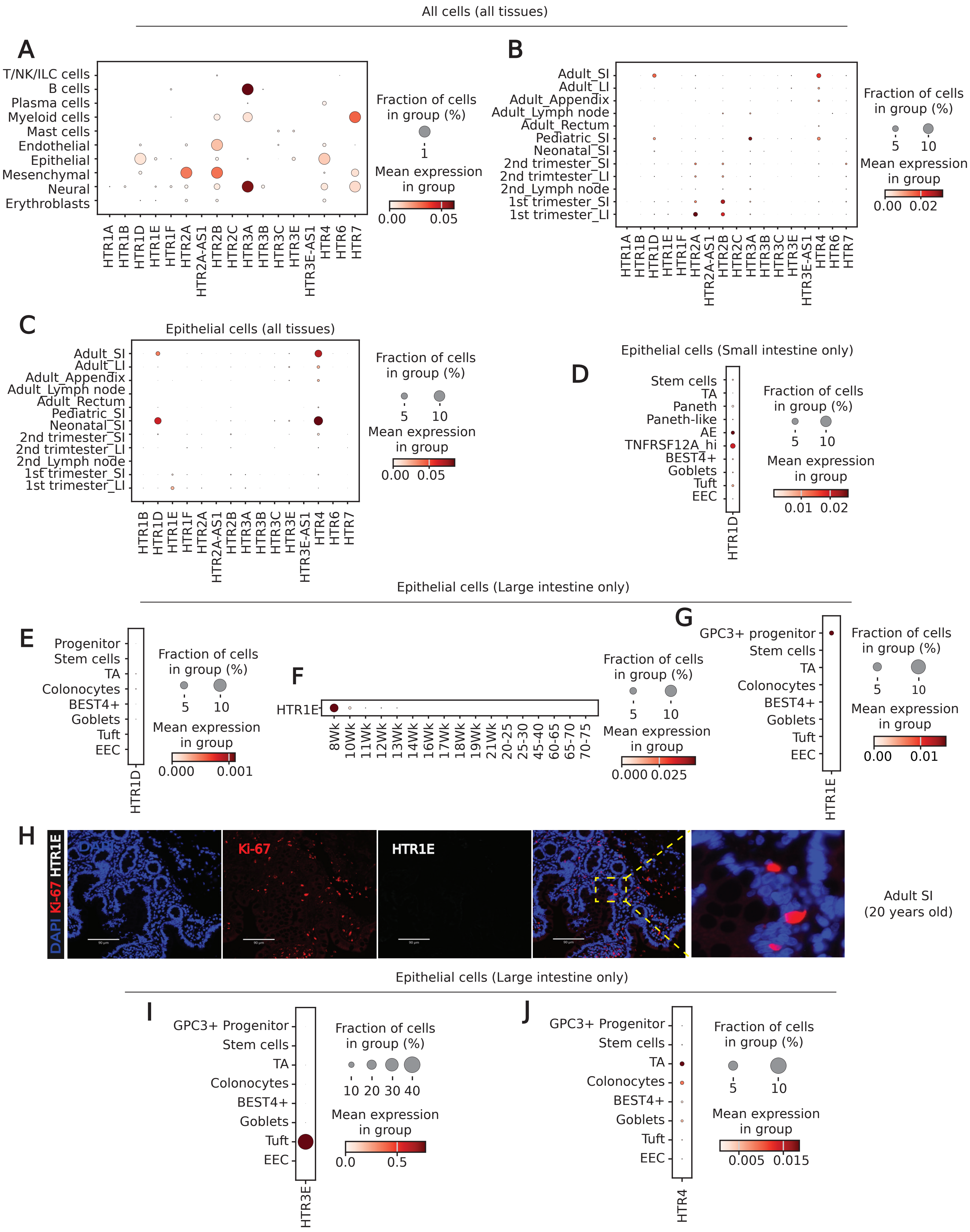
